# Species-specific interactions with apex carnivores yield unique benefits and burdens for mesocarnivores

**DOI:** 10.1101/2024.08.17.608414

**Authors:** Wesley Binder, Jack W. Rabe, Zoe Lowe, Gordon Scott, Claire Lacey, Eliza King, Daniel R. Stahler, Stotra Chakrabarti

## Abstract

Mesocarnivores navigate a complex *risk-reward* continuum in ecosystems shared with their apex counterparts, balancing scavenging opportunities with risks of mortality. However, the risks to mesocarnivores in multi-carnivore systems are not uniform—often varying with specific apex-meso pairings. Using camera traps and GPS-telemetry, we examined space-use, temporal activity, fine-scale interactions, and scavenging behavior of apex (wolves, cougars) and mesocarnivores (red foxes, coyotes) in northern Yellowstone National Park, USA. Coyote space use was positively linked to wolf presence, while red fox space use was positively linked to cougar presence. Notably, photo-detection rates of mesocarnivores doubled within 24 hours of apex carnivore detections—except for coyotes following cougars. Coyotes were more frequent scavengers at wolf and cougar kills than red foxes, but they also suffered the highest mortality rates. Over 60% of wolf caused coyote mortalities were linked to wolf kill-sites, though wolves rarely consumed the coyotes themselves. In contrast, cougars hunted and consumed coyotes as prey, posing an additional risk of predation. These findings highlight how divergent hunting strategies and habitat preferences between canid and felid apex carnivores create separate risks for mesocarnivores. Our study offers new insights into the species-specific behavioral dynamics that can shape trophic interactions in multi-carnivore systems.

## INTRODUCTION

Intraguild carnivore interactions play a pivotal role in ecosystem function, shaping the abundance, distribution, and behavior of both carnivores and their prey through top-down effects [1]. These interactions, shaped by the complex strategies employed by carnivores across different trophic levels to coexist, are particularly important to understand given that human activities have disproportionately disrupted apex carnivores in the Anthropocene. Humans have both driven significant population declines and local extinctions of apex carnivores in some areas [1] and facilitated their recovery through reintroduction and conservation efforts in others [2–4].

These actions, in turn, impact carnivore communities, as apex carnivores can affect mesocarnivores both directly and indirectly. For instance, wolves (*Canis lupus*) can suppress coyote (*Canis latrans*) populations through both lethal removal and resource competition, which in turn reduces the top-down effects that coyotes exert on red fox (*Vulpes vulpes*) populations [5]. Thus, the extent to which mesocarnivores are affected by the presence of apex carnivores depends on their relative position within the trophic community.

However, the factors mediating the dynamics between apex and mesocarnivores are complex and often difficult to untangle. While apex carnivores can suppress mesocarnivore populations, they can also benefit them by providing scavenging opportunities from the carcasses of their prey [6–8]. As a result, mesocarnivore distributions and behavior in relation to their apex counterparts depend on a delicate balance of risks (interference competition) and rewards (carcass availability) that vary in both space and time [9].For instance, a global review revealed that 30% of mesocarnivore diets were composed of scavenged ungulate carcasses, yet a nearly identical proportion—about one-third—of their deaths were attributed to apex carnivores [10].

Larger mesocarnivores, in particular, were more reliant on scavenging opportunities, and this facilitation appeared to be more costly in less productive environments, leading to greater suppression from apex carnivores [10]. The relationships between mesocarnivores and apex carnivores are thus shaped by several factors, including the mesocarnivores’ reliance on scavenging to supplement their diets, their ability to locate apex carnivore kill sites quickly, and the extent to which apex carnivores influence mesocarnivore mortality risks.

The risks faced by mesocarnivores also depend on how they interact with specific apex carnivores. For example, when mesocarnivores are primarily competing for resources—such as carcasses—their risks mostly arise from direct encounters with apex carnivores at kill sites [10]. In contrast, if mesocarnivores are killed by apex carnivores independent from carcass sites, then their risks are influenced by the habitat preferences and activity patterns of the apex carnivore [11]. Such relationships likely drive significant variation in intraguild mortality rates across carnivore family groups. For example, canid mesocarnivores experience mortality from canid apex carnivores at rates five times greater than from felid apex carnivores [10]. Thus, the risks and rewards mesocarnivores experience when associating with their apex counterparts should vary with specific meso-apex pairings, yet the mechanisms driving such differences are poorly understood. Such species-specific interactions can be critical in further mediating mesocarnivore *landscapes of fear*, and in turn have effects on entire ecosystems.

In the Northern Range of Yellowstone National Park, wolves and cougars (*Puma concolor*) are abundant during winter when they are supported by a diverse assemblage of herbivores such as elk (*Cervus canadensis*), bison (*Bison bison*) and deer (*Odocoileus* spp.) that congregate at high densities [12–14]. Wolves and cougars kill larger ungulate prey during these months [15–17], and such large carcasses provide scavenging opportunities to a host of mesocarnivores including coyotes and red foxes [8]. Such scavenging opportunities appear to be critical for mesocarnivores in the winter because much of their regular prey (small mammals and ungulate neonates, birds, fruits, insects) become scarce and/or subnivean [8]. Coyotes and red foxes, however, also navigate the threat of lethal encounters with wolves and cougars while accessing these carcasses, and in general. To examine how competitive and facilitative relationships vary by specific apex-mesocarnivore pairings, we evaluated the degree to which coyotes and red foxes exhibited spatial and temporal avoidance, or attraction, to wolves and cougars in this system.

We expected each apex-mesocarnivore pairing to exhibit distinct associations for two reasons. First, long-term anecdotes from Yellowstone suggest the mechanisms of lethal interactions for mesocarnivores differ by apex carnivore species. Cougars appear to hunt mesocarnivores as prey, while wolves are more likely to kill them at ungulate-carcass sites as a form of resource defense. Therefore, we anticipate mesocarnivores to exhibit more avoidance of cougars compared to wolves given the higher threats associated with cougar predation. Second, we expected coyotes to exhibit more attraction to apex carnivores than red foxes as their greater size and energetic demands should increase their reliance on scavenging kills when alternative resources are scarce. To explore these relationships, we conducted winter camera-trapping surveys to assess occupancy, diel activity patterns, and fine-scale spatiotemporal responses of apex and mesocarnivores. We also monitored predation events of GPS-collared wolves and cougars to quantify mesocarnivore scavenging at kills, as well as direct killings of mesocarnivores.

## MATERIALS AND METHODS

### Study Area

We conducted the study in the Northern Range of Yellowstone National Park (Figure 1), an area with elevations ranging from 1500 to 2400 meters. The landscape features a mix of rugged, rocky terrain and open, rolling hills. Common vegetation types in the region include Douglas-fir (*Pseudotsuga menziesii*), sagebrush (*Artemisia* spp.), and juniper (*Juniperus spp.*). The area experiences long, cold winters, with average snow-water equivalents ranging from 2 to 30 cm [18]. Apex carnivores, such as wolves, grizzly bears (*Ursus arctos horribilis*), black bears (*U. americanus*), and cougars, are present in relatively high numbers, while coyotes, red foxes, bobcats, and martens (*Martes americana*) are the prominent mesocarnivores [14,17].

**Figure 1.**
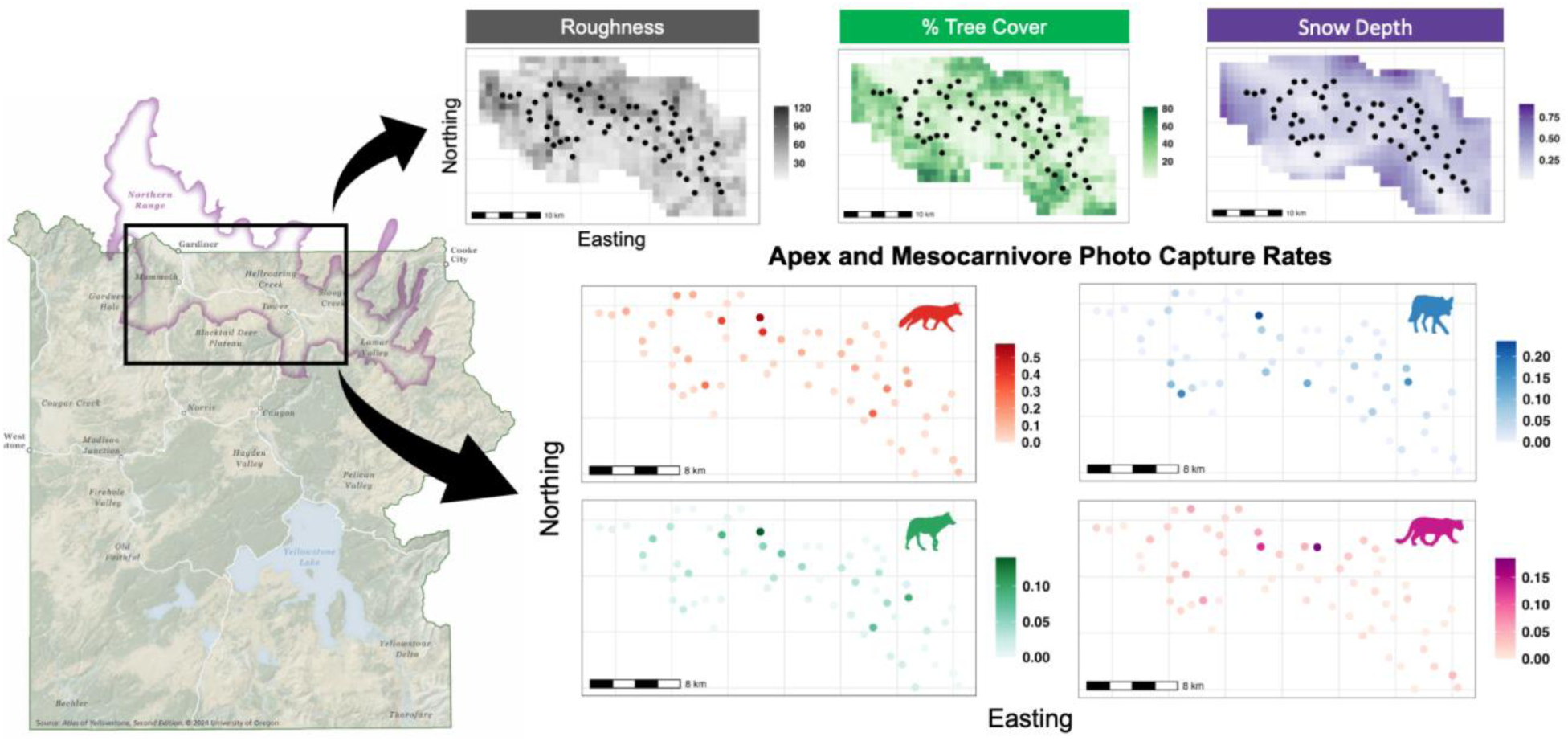
Locations of camera stations, gradient of habitat covariates, and photo-capture rates of mesocarnivores (red fox = red, coyote = blue) and apex carnivores (wolf = green, cougar = pink) in northern Yellowstone National Park, USA.

### Camera Trap Surveys

Camera surveys were conducted within a 580 km² area, defined by a 5-km buffer surrounding our camera-trap grid, which was designed to estimate cougar densities. We deployed camera stations in a “checkerboard” pattern to increase the variability in station spacing [19], using a grid of 3.2 km² cells [20]. In each selected grid cell, we set up two camera stations at locations chosen to maximize detections, such as natural movement corridors or pinch points, with a minimum distance of 1 km between stations.

Camera stations were active during 15-week winter sampling periods (referred to as “sessions”) from December to March in the years 2020-21, 2021-22, and 2022-23 (hereafter referred to as 2021, 2022, and 2023, respectively). We used Browning Spec Ops trail cameras equipped with motion-activated passive infrared sensors. In 2021, we deployed 90 cameras across 60 stations, with half of the stations having a single camera and the other half having two cameras placed opposite each other. For 2022 and 2023, all stations were equipped with two cameras, bringing the total number of cameras to 120. Each camera was configured for long-range motion detection and recorded 20-second videos with a one-minute delay between recordings. We manually reviewed all videos, identified species, and extracted metadata using the Timelapse2 image analysis software [21].

### Data Analysis

#### Multi-Species Occupancy Models

To evaluate how habitat covariates and spatial interactions with other carnivores influence the occurrence of apex and mesocarnivores, we constructed multi-species occupancy models [22] using the *unmarked R* package [23,24]. Occupancy models incorporate binary presence-absence (1/0) data through likelihood-based approaches to estimate species occurrence at site *i* in a latent *z* (occupancy) “state” model, while accounting for imperfect detections through a separate “detection” model. Multi-species occupancy models are an extension of the single-species occupancy model [25] that apply a hierarchical modeling framework to estimate the effect of covariates (first-order) and species co-occurrence (second-order) on occupancy.

We used 1-km^2^ grid cells for modeling occupancy, with each cell containing no more than one camera station. Because wolves, cougars, coyotes, and potentially red foxes typically occupy ranges larger than 1 km^2^, the close spacing of our cameras may violate assumptions of spatial independence. Hence, our occupancy estimates more aptly describe patterns of “space use” rather than distribution [26]. For estimating detection probabilities of each species, we considered two covariates: the number of active cameras at a station and snow depth. As the number of active cameras at a station (ranging from 0–2 as a product of how many cameras were active and for how long during week-long occasions) increased, we expected an increase in detection probability. We used mean snow depth at camera stations during each week-long occasion at a 1-km^2^ resolution using SNODAS [27]. We expected a negative relationship between snow depth and detection probability, such that increasing snow depth as winter sessions progressed would lead to fewer individuals moving through camera stations, lowering detection probability. For estimating species occupancy, we included topographic roughness, percent tree cover, and average snow depth at a 1-km^2^ spatial resolution to account for bottom-up processes, such as habitat and landscape characteristics, that influence carnivore distributions [28]. These habitat covariates were included because they are known to affect the occupancy and movement of carnivores [14]. Please refer to Table S1 for covariate details.

Our primary interest was in estimating species co-occurrence patterns and the effects of landscape covariates on occupancy, rather than temporal occupancy dynamics. We therefore adopted a ‘stacked’ data structure, where each site ⨉ year combination was treated as an independent site [29,30], resulting in 178 sites (58 in 2021, and 60 in 2022 and 2023). To account for the non-independence of sites across years, we included year as a fixed effect on both detection and occupancy in all models. Using a two-stage model selection approach [31], we first constructed single-species models to evaluate which covariates best explained detection probability while holding occupancy constant at the null model (*Ψ*(.)). For each species, we used Akaike’s Information Criterion (AIC) to select the best detection model from four candidate models representing all possible additive combinations of the two detection covariates, including the null model with only year as a fixed effect. Second, we carried over the top detection sub model to select the best first-order model by considering all possible additive combinations of covariates on occupancy (eight candidate models for each species). Following Arnold (2010), we accounted for parameter redundancy by selecting more parsimonious models if they were within 2 AIC units of a model with more parameters [32]. After selecting the top first-order model, we added second-order species interactions to develop multi-species models that tested the effects of species co-occurrence on occupancy. Please refer to Table S3 and S4 for a list and comparison of candidate models.

#### Temporal Activity Patterns

We used the *activity* package in R to estimate diel activity patterns for wolves, cougars, coyotes, and red foxes using non-parametric kernel density estimation [33]. To maintain independence of detections, photo-captures of the same species within 30 minutes were omitted [34]. To quantify temporal niche partitioning between species and identify potential coexistence strategies that extend beyond broad-scale occupancy, we used the *overlap* package in R to assess overlap of diel activity patterns between each apex-meso pairing by calculating the area under the combined activity curves, [inline] [35]. We generated 10,000 bootstrap samples of the activity overlap coefficient, [inline], for each species-pair to estimate 95% confidence intervals of overlap.

#### Time-to-event

We examined fine-scale mesocarnivore activity in response to apex carnivore movements using time-to-event analysis [36,37]. The elapsed time between a mesocarnivore detection following an apex carnivore detection was recorded at each camera station for a given session.

Subsequently, we combined data across all sessions for each apex-meso pairing. We truncated apex carnivore detections to only include events that were followed by a mesocarnivore detection (i.e., apex carnivore detections followed by another apex carnivore detection were omitted). Detections of mesocarnivores were recorded for 5 days following an apex carnivore detection, with elapsed time differences binned into 24-hour periods. We estimated detection probabilities by dividing the number of mesocarnivore detections in each 24-hour period by the total number of detections across all sessions [36,37]. These detection probabilities were compared to random iterations (n = 1,000) of mesocarnivore detections to derive expected detection probabilities for each 24-hour period by using standard two-tailed permutation tests [37]. If mesocarnivores were avoiding apex carnivores through fine-scale temporal segregation , we would expect coyote and red fox detection probabilities to be significantly lower following an apex carnivore detection. The opposite would apply if they were instead tracking/following wolves and cougars, likely to acquire food by scavenging apex carnivore kills .

#### Monitoring of Apex Carnivore Predation Events

To assess mesocarnivore scavenging at apex carnivore kills and direct killing of mesocarnivores, we monitored predation by GPS-collared wolves (n = 26 from 8 packs) and cougars (n = 17) during 30-day winter sampling periods (November to March) from 2016–2023. Monitored wolves and cougars were fitted with Vectronics or Telonics GPS-Satellite collars that were programmed to collect hourly fixes. We searched GPS-location clusters [38] to identify kill sites and conducted necropsies on prey remains. At each kill, we ascertained the likely cause of death, the species, sex, age of prey when possible, and the presence of scavengers through visual observations and sign (e.g., scat, tracks, hair). To complement our camera-trap analysis, we also filtered this long-term predation data by selecting scavenging and predation events that occurred only during the three winter camera-trap sessions, and assessed whether the trends during those years differed from our longer-term data (Figure S5).

## RESULTS

Across all three 15-week winter camera-trapping sessions, we collectively monitored for 18,617 sampling days. Red foxes were detected at 88% of sites, coyotes at 56% of sites, wolves at 53% of sites, and cougars at 47% of sites (Figure S2).

### Multi-Species Occupancy Models

#### Detection probability

Variation in red fox detection probability was best explained by snow depth, with red fox detection increasing as snow depth decreased (*β* = -0.21, 95% CI = -0.34 to -0.08, *P* = 0.0018). Holding snow depth at its mean, red fox detection varied by year with marginal detection probabilities of 0.33 (95% CI = 0.29 – 0.36) in 2021, 0.24 (95% CI = 0.21 – 0.28) in 2022, and 0.39 (95% CI = 0.34 – 0.43) in 2023. While neither the number of active cameras nor snow depth significantly affected coyote or wolf detection probability, there was variation between sessions. Coyote detection probability was 0.25 (95% CI = 0.21 – 0.28) in 2021, 0.14 (95% CI = 0.11 – 0.18) in 2022, and 0.22 (95% CI = 0.18 – 0.27) in 2023. Wolf detection probability was 0.17 (95% CI = 0.14 – 0.21) in 2021, 0.12 (95% CI = 0.09 – 0.16) in 2022, and 0.21 (95% CI = 0.18 – 0.26) in 2023. Cougar detection probability did not vary annually, but did increase significantly as snow depth decreased (*β* = -0.27, 95% CI = -0.51 to -0.03, *P* = 0.03).

#### Occupancy probability

In the top-performing first-order model, habitat covariates only explained significant variation in cougar occupancy, while the null “year-only” model performed best for red fox, coyote, and wolf. Cougar occupancy probability increased as mean snow depth decreased, (*β* = - 1.06, 95% CI = -1.99 to -0.13, *P* = 0.025) and roughness increased (*β* = 1.4, 95% CI = 0.78 – 2.03, *P* < 0.0001; Figure S2). After incorporating second-order species interactions on occupancy, we found that coyote occupancy was positively associated with wolf occupancy (*β* = 1.62, 95% CI = 0.81 – 2.42, *P* < 0.0001). Coyote occupancy was 0.75 (95% CI = 0.58 – 0.86) at sites where wolves were present and 0.35 (95% CI = 0.20 – 0.55) where wolves were absent (Figure 2b). Red fox occupancy was positively associated with cougar occupancy (*β* = 1.94, 95% CI = 0.51 – 3.36, *P* = 0.0077) (Figure 2b). Where cougars occurred, red fox occupancy was 0.96 (95% CI = 0.86 – 0.99), compared to 0.82 (95% CI = 0.63 – 0.91) where cougars did not occur.

**Figure 2.**
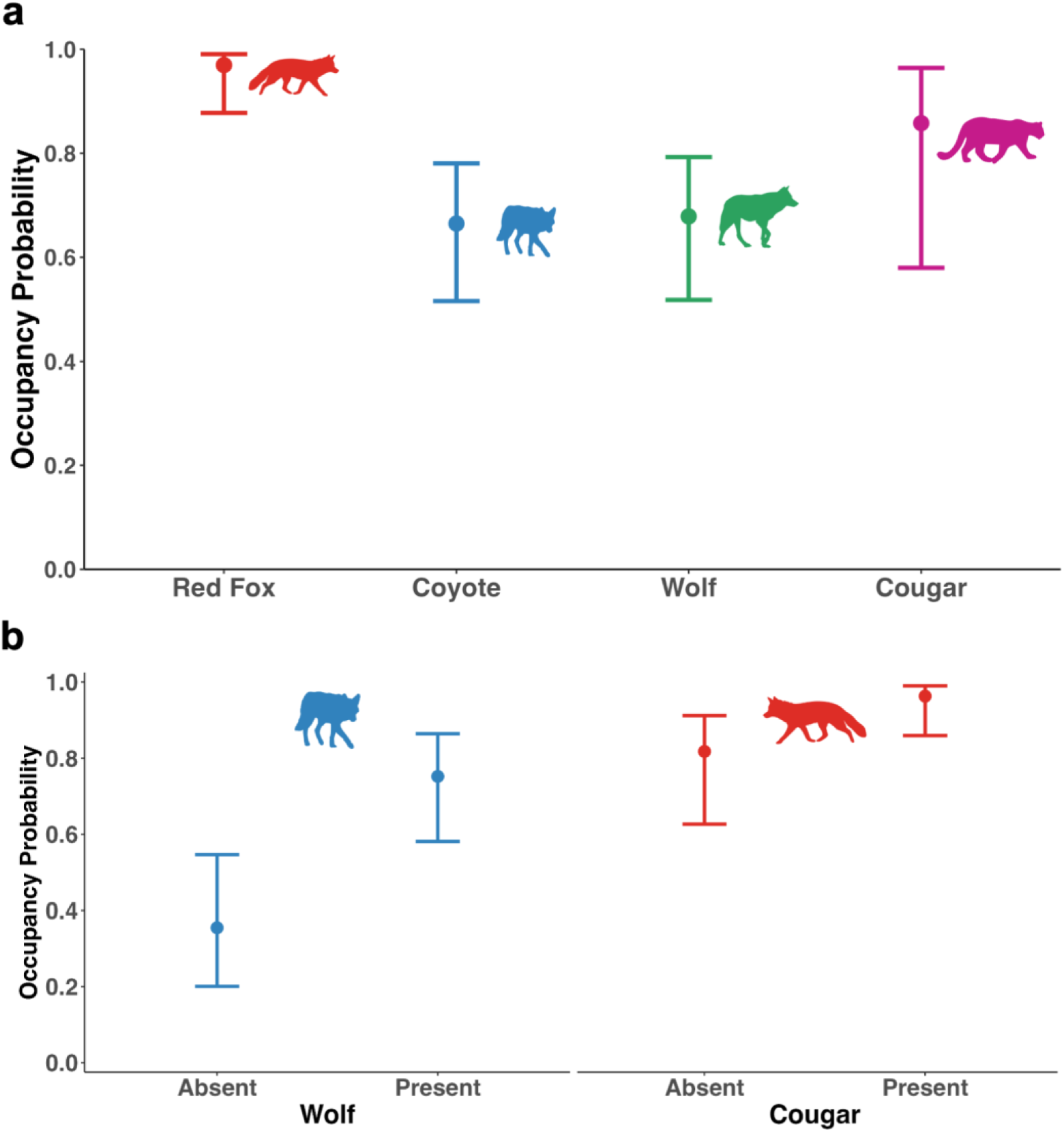
**(a)** Apex and mesocarnivore marginal occupancy probabilities estimated from camera trapping surveys between 2021–2023. **(b)** Coyote occupancy probability for 2021–2023, conditional on wolf presence-absence and red fox occupancy probability for 2021–2023, conditional on cougar presence-absence. Occupancy probabilities were based on the best-supported, second-order multi-species occupancy model.

### Temporal Activity Patterns

Cougars (n = 341 detections) exhibited unimodal diel activity, being most active around dusk peaking at 1800 and least active from morning (0800) to midday (1400) (Figure 3). Wolves (n = 443 detections) had bimodal, crepuscular activity patterns, with peak activity shortly after dawn (∼0800–0900) and a secondary peak at dusk (1700–1800) (Figure 3). Coyotes (n = 541 detections) had more consistent activity across the diel cycle and were more diurnal, with peak activity occurring from late morning (0900) to midday (1400) and the lowest activity before dawn (0500) and after dusk (2100) (Figure 3). Red foxes (n = 1520 detections), on the other hand, exhibited highly nocturnal patterns, being most active shortly after dusk (1800–2100) and sustaining high activity through the night (2100–0600) with little activity during daylight hours (0800–1600) (Figure 3). Temporal overlap between apex and mesocarnivores was 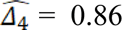 (CI: 0.81 – 0.89) for wolf-coyote, 0.68 (CI: 0.61 – 0.7) for wolf-red fox, 0.81 (CI: 0.74 – 0.85) for cougar-coyote, and 0.72 (CI: 0.66 – 0.76) for cougar-red fox across the three sessions (Figure 3a). Overlap between mesocarnivores (red fox-coyote) was 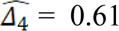 (CI: 0.53 – 0.62), while overlap between apex carnivores (wolf-cougar) was 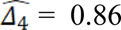 (CI: 0.8 – 0.91) (Figure 3b).

**Figure 3.**
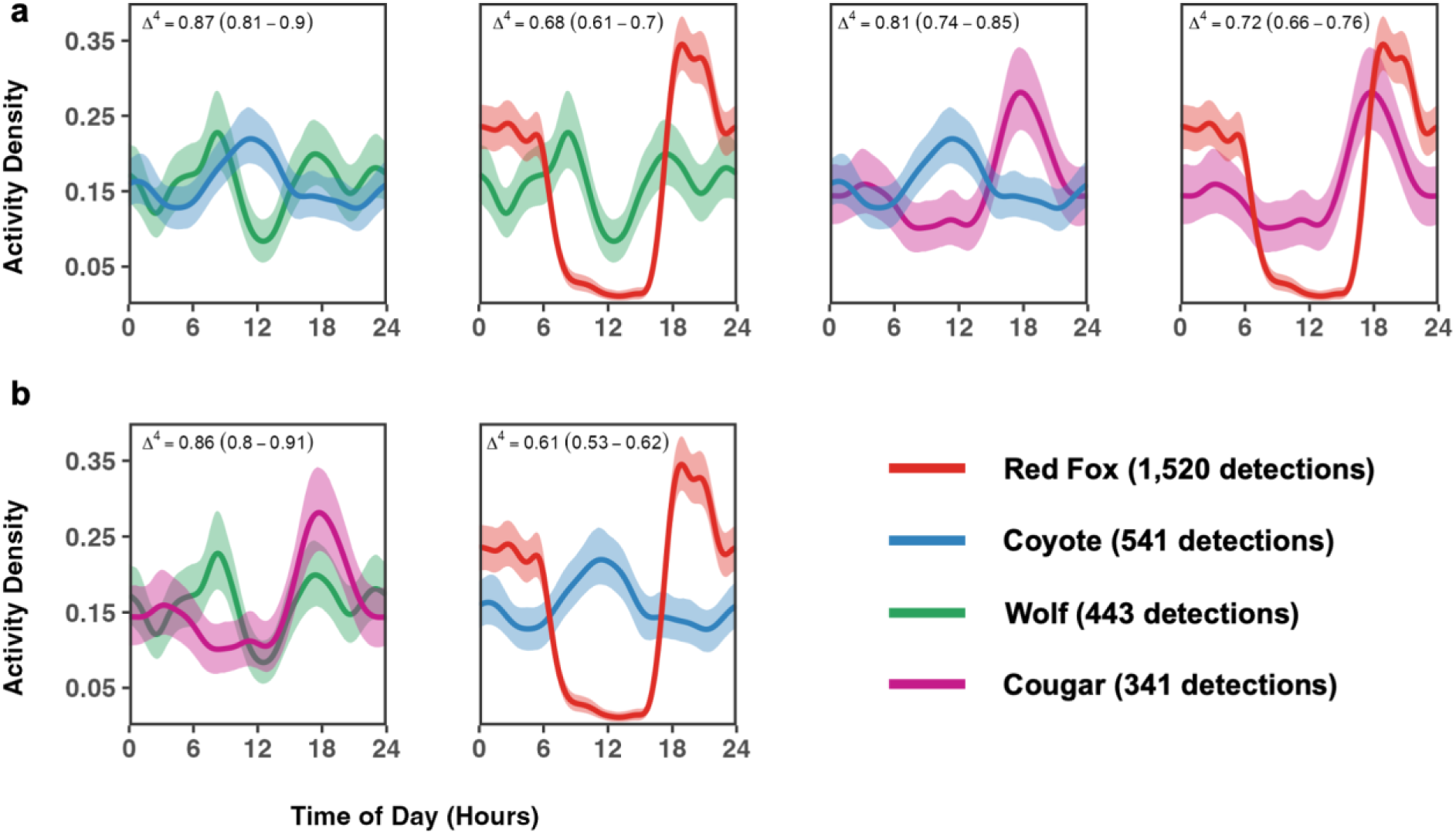
Diel activity patterns with 95% bootstrapped CIs for study carnivores. Mean activity overlap **(a)** between apex and mesocarnivores, and **(b)** between apex carnivores and between mesocarnivores.

### Time-to-event

Compared to the mean values derived from random iterations of detection probabilities, red foxes and coyotes were both over twice as likely to be detected in the 24-hour period following a wolf detection (red foxes: 0.034 compared to 0.015, *P* = 0.001; coyotes: 0.05 compared to 0.018; *P* = 0.001; Figure 4a). However, their detection probabilities following cougar detections differed. While red foxes were over twice as likely to be detected in the 24- hour period following a cougar detection (0.024 compared to 0.011; *P* = 0.001), coyote detection rates were not affected. Additionally, red foxes were more likely to be detected in the 72-hour period following a wolf detection (0.01 compared to 0.007; *P* = 0.04) and the 96-hour period following a cougar detection (0.009 compared to 0.004; *P* = 0.001). Overall, the time interval between detections of a mesocarnivore subsequent to an apex carnivore detection was shortest for wolf-coyote (median = 17.67 hours) and longest for cougar-coyote (median = 57.35 hours). Foxes showed similar latency to wolf and cougar detections, with medians of 21.11 and 24.69 hours, respectively (Figure 4b).

**Figure 4.**
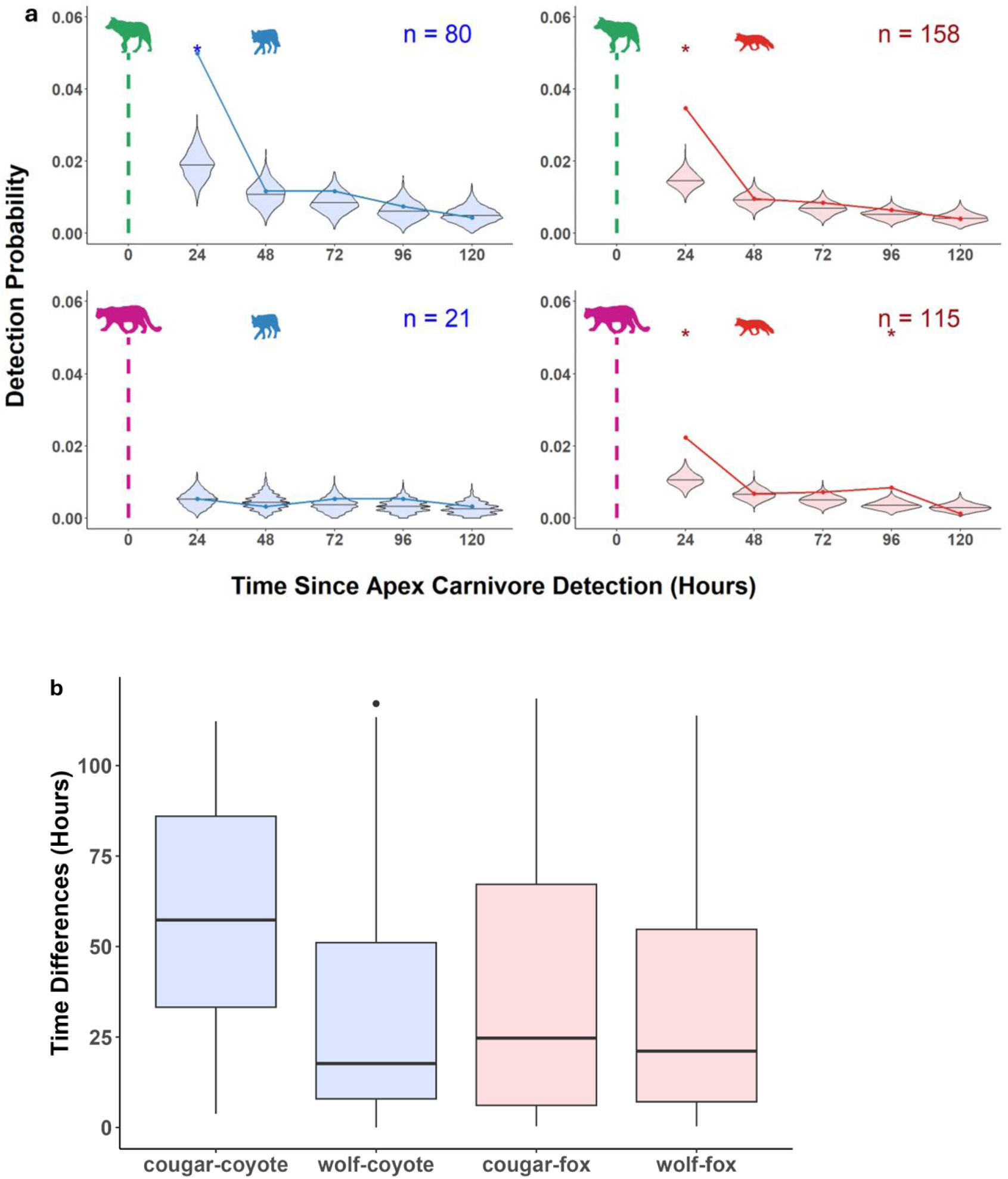
**(a)** Daily binned temporal attraction-avoidance of mesocarnivores in response to apex carnivore detections at camera stations. Apex-mesocarnivore pairs include wolf-coyote (top-left), wolf-red fox (top-right), cougar-coyote (bottom-left), and cougar-red fox (bottom-right). Points and lines represent observed mesocarnivore detection probabilities, while violin plots show the distribution of expected mesocarnivore detection probabilities based on random movements (i.e., no attraction or avoidance). *denotes significant differences between observed and expected detection probabilities. **(b)** Median time differences of mesocarnivore detections following apex carnivore detections.

### Monitoring of Apex Carnivore Predation Events

We detected 327 wolf kills and 257 cougar kills through winter predation monitoring (2016–2023). Coyote presence was recorded at 222 (68%) wolf kills and 80 (31%) cougar kills (Figure 5a). Red fox presence was recorded at 67 (20%) wolf kills and 46 (18%) cougar kills (Figure 5a). From this predation monitoring, we documented 18 coyotes and one red fox killed by wolves (Figure 5b). Of the coyotes killed by wolves, 11 (61%) occurred at wolf feeding sites (i.e., ungulates killed or scavenged by wolves), while the red fox mortality did not occur at a wolf-killed prey site. Cougars killed eight coyotes and three red foxes during this time (Figure 5b). None of these mortalities were associated with cougar-killed prey sites, rather they were fully consumed and generally found in cache piles - suggestive of cougar consumption. We found no major differences between the longer-term data and the data overlapping with our camera-trapping study (see Figure S5 for filtered results).

**Figure 5.**
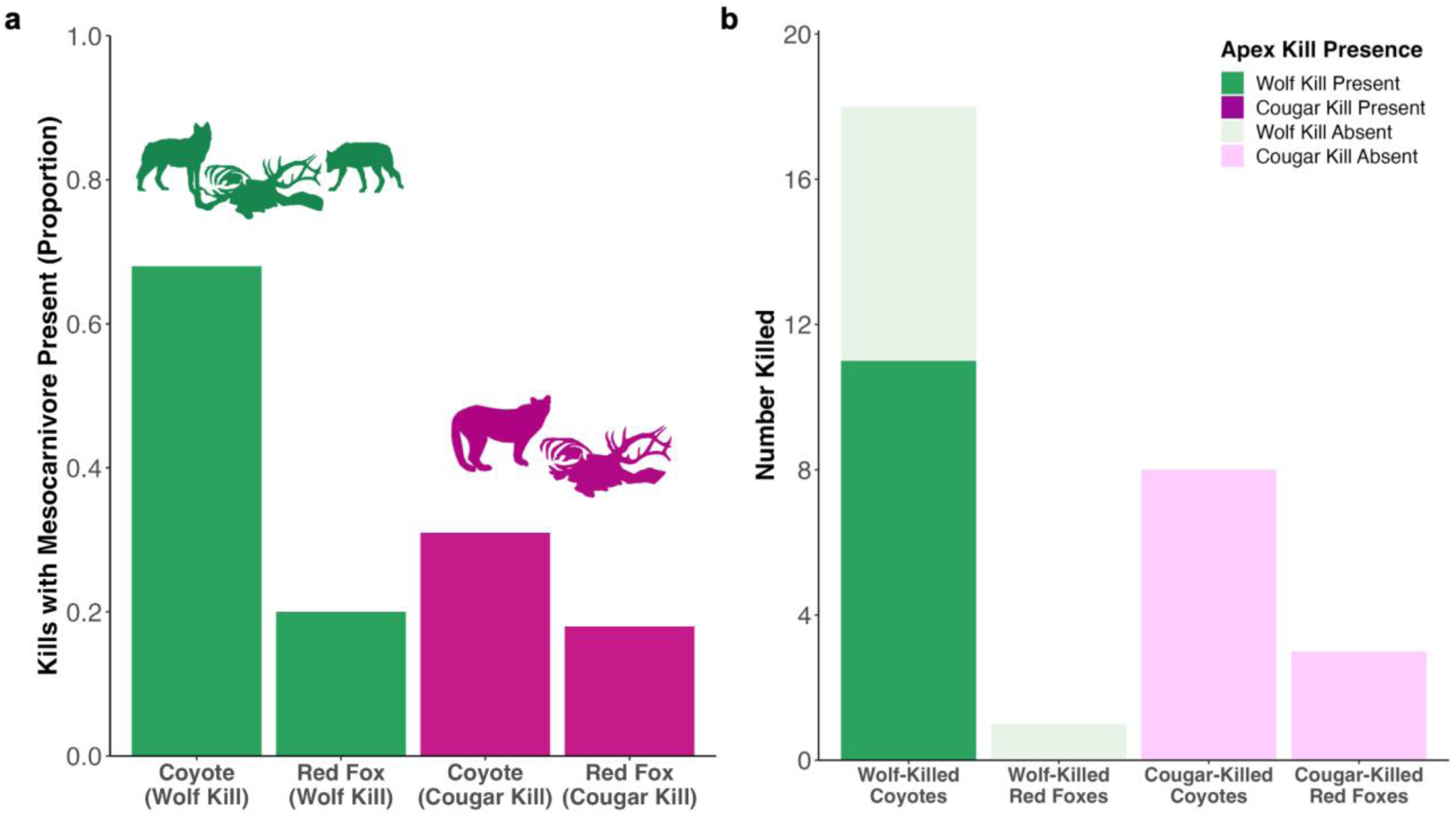
Wolf (n = 327) and cougar (n = 257) kills, surveyed through GPS-cluster searches of collared wolves and cougars between 2016-2023, bearing **a)** presence of coyote and red fox scavenging expressed as a proportion, and **b**) counts of coyote and red fox killed. For methods see main text. Wolf and elk carcass graphics by Kira Cassidy.

**Figure 6.**
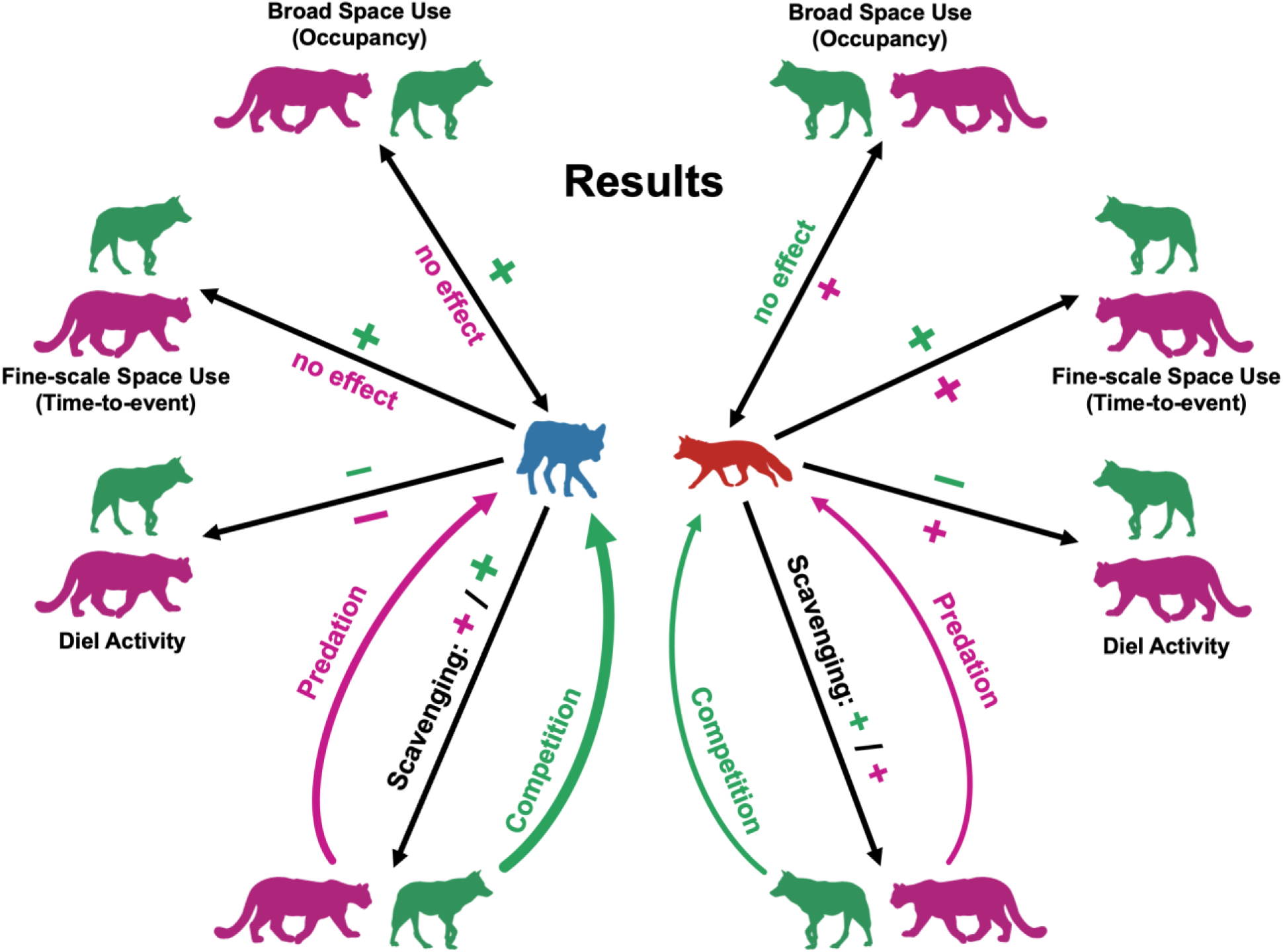
Conceptual representation of results from analyses for each apex-mesocarnivore pairing. Mesocarnivores (Coyote = blue, Red fox = red) are shown in the center and apex carnivores around the perimeter (Wolf = Green, Cougar = Pink). For “Occupancy/ Broad Space Use,” the multi-directional arrow refers to co-occurrence results from the top second-order multi-species occupancy model. “Scavenging” represents the proportion of ungulate kills made by an apex carnivore that were visited by the respective mesocarnivore. “+” represents a positive association/effect, “–” represents a negative association/effect, and “no effect” indicates an insignificant result. Effects are colored green or pink to indicate they are affiliated with wolf or cougar results, respectively. The size of the “+” or “–” is positively-related to the magnitude of the effect observed.

## DISCUSSION

Ecological communities are influenced by the top-down effects that carnivores exert on prey populations, but these effects also extend to other carnivores [1]. Intraguild carnivore interactions are complex in that the negative impacts of competition can be partially offset by the scavenging opportunities competitors provide with their kills. While these tradeoffs are common between apex and mesocarnivores, we found considerable variation across species-specific pairings.

Red foxes and coyotes were both ubiquitous in our study area and common scavengers at wolf and cougar kills, but exhibited unique spatial and temporal relationships with each apex carnivore. Our multi-species occupancy models revealed positive associations between the space use of coyotes and wolves, as well as red foxes and cougars. Further, red foxes and coyotes exhibited opposite diel activity patterns (Figure 3b), as the nocturnal activity of red foxes closely aligned with that of cougars, while coyotes were most active mid-day when apex carnivores were least active. Their fine-scale movements also differed in response to apex carnivore presence.

Red fox detection probabilities at camera stations doubled in the 24 hours following a wolf or cougar detection, but coyotes only displayed this attraction to wolves, not cougars (Figure 4a). Accordingly, we found that coyotes were more frequent scavengers at wolf kills compared to cougar kills. However, scavenging on wolf kills was often costly, as over half the coyote mortalities attributed to wolves during our study occurred at wolf kill sites. Interestingly, we did not document wolves consuming the coyotes they killed, likely indicating that such lethal actions resulted out of resource defense. Conversely, all coyotes killed by cougars occurred as standalone predation events (i.e., independent of ungulate carcass sites) in which cougars consumed the coyotes. The ambush hunting style of cougars, combined with the forested and topographically rough terrain they inhabit, could lead to coyotes being more vulnerable in such habitats [39]. In contrast, the open and flatter terrain preferred by wolves could provide more visual and olfactory cues for coyotes to assess (and escape) immediate risks of lethal encounters. Such disparities in apex carnivore hunting strategies seemingly tie back to the spatiotemporal attraction of coyotes towards wolves but not for cougars, and highlight the nuances of such competitive outcomes.

As mesocarnivores adapt their coexistence strategies to different apex carnivore species, their behaviors are likely to shift more dramatically when multiple competitors are present. For example, high winter apex carnivore densities in our study system may explain the midday peak in activity for coyotes, which differed from the more crepuscular activity patterns of coyotes inhabiting adjacent areas of the Greater Yellowstone Ecosystem with lower apex carnivore densities [40]. In contrast, red foxes appeared less sensitive to the diel activity of apex carnivores. Not only did their peak in activity align with that of cougars, but they also displayed the same nocturnal patterns as red foxes across the Greater Yellowstone Ecosystem [40].

Notably, red foxes exhibited high temporal segregation from coyotes, having the lowest temporal overlap of any species pair (Figure 4). Studies elsewhere have shown that coyotes and red foxes compete severely, with coyotes competitively excluding and/or killing red foxes [5,41]. Thus, the nocturnal activity of red foxes could be the result of avoiding peak coyote activity during the day that is in turn mediated by wolf and cougar activity (Figure 4). Temporal sympatry amongst the carnivore guild appears to be layered, with apex carnivores affecting coyotes, and coyotes affecting red foxes.

Apex- and mesocarnivore relationships depend on competition that occurs within trophic groups, as spatial and/or temporal responses to such competitors alters the availability of ecological niche space for other carnivores in the community. For example, cougars in Yellowstone changed their winter space use after wolves were reintroduced by increasing their selection for areas with escape terrain (e.g., ruggedness, tree cover) [17]. This avoidance of wolves could in turn facilitate coyote space use, providing habitats where encounters with cougars are scarce. Our results support this, as coyote space use, fine-scale movements, and scavenging rates were more strongly associated with wolves than cougars. Accordingly, if red foxes experience the most severe competition from coyotes, then greater overlap with cougar space use could alleviate such competition. While red foxes scavenged wolf and cougar kills at a similar rate, our analyses revealed substantial overlap between cougar and red fox space use and activity patterns. Mortalities from cougars still occurred, but we believe that foxes are more adept at avoiding immediate risks of cougar predation during encounters through their increased agility, as well as their relatively large foot:body ratio that allow for better maneuverability on top of snow [42]. Coyotes being heavier, might not be as lucky when encountering cougars. Thus, competition within trophic groups likely plays a key role in shaping the apex-mesocarnivore interactions we observed, creating scenarios akin to “the enemy of my enemy is my friend”.

In contrast, competition between apex- and mesocarnivores is often driven by scavenging, where apex carnivores take measures to reduce food loss (e.g., carcass defense, caching, etc.) and mesocarnivores balance the need for such dietary subsidies with the associated risks of confrontations at carcass sites [10]. Thus, the availability of alternative resources for mesocarnivores should influence the extent to which they seek out scavenging opportunities and the overall strength of their relationship with apex carnivores. In northern Yellowstone, the limitations of prey acquisition during winter months may increase mesocarnivore reliance on scavenging, particularly for coyotes who appear to depend on these subsidies more so than smaller red foxes (Figure 5) [40,43]. Foxes may be better suited to meet their energetic demands by hunting themselves, as they are commonly observed hunting subnivean rodents by pouncing through the snowpack during the winter (D. Stahler *pers.comm.*). Therefore, apex carnivores facilitate coyotes and red foxes differently, and these relationships are likely to change during the spring/summer when the availability of rodents, neonatal ungulates, vegetation, and insects increase [43]. The presence of grizzly and black bears that emerge in the summer after winter-long hibernations can also complicate these relationships because bears often limit carrion consumption for other scavengers [44].

While our study highlights the positive aspects of apex and mesocarnivore interactions through scavenging, we did not explore variations in red fox and coyote population trends. In fact, previous research documented a 39% decline in coyote abundance following wolf reintroduction in the Lamar Valley of Yellowstone, an area adjacent to our study area [45].

Additionally, Merkle et al. (2009) showed a decrease in wolf-coyote interactions over time (between 1997 and 2007) in our study area, indicative of changes to coyote density and/or behavior (e.g., wolf avoidance) [46]. Despite benefiting from carrion made available by wolves and cougars, red fox and coyote populations may still experience top-down population regulation via direct killing that maintains them at low densities [47]. Long-term studies that estimate population trends of mesocarnivores with respect to apex carnivore densities are required to test such effects.

The magnitude and impact of trophic-mediated responses related to large carnivore recoveries have become a contentious issue, perhaps nowhere more so than in Yellowstone National Park [48–50]. Although the reintroduction of wolves and recolonization of cougars to this area has likely altered mesocarnivore distributions and abundance, red foxes and coyotes also benefit from living with their dominant counterparts. Overall, red foxes and coyotes in our study area exhibited more attraction than avoidance to apex carnivores during winter months, when small prey and alternative food resources are scant. Such attraction, possibly a consequence of desperate foraging decisions in winter, can also prove fatal. However, risks to mesocarnivores are uneven across dominant-subordinate carnivore pairings, with apex carnivore behavior changing the drivers of these interactions, adding nuance to the *landscape of fear* for mesocarnivores.

## Acknowledgements

We are grateful to the numerous field technicians from the Yellowstone Wolf and Cougar Projects who made data collection possible. We would also like to thank the fStop Foundation for providing necessary equipment. We acknowledge Yellowstone Forever (and their contributing donors), the National Science Foundation, Oregon State University, University of Minnesota, National Park Service, US Geological Survey, and Macalester College for funding support.

## SUPPLEMENTARY INFORMATION

**Table S1:** Covariate List for Detection and Occupancy Models & their Description and sources.

| Occupancy Covariates |  |  |
| --- | --- | --- |
| Variable | Type, Range | Description |
| Cover | Continuous | Values accrued from the Rangeland Analysis Platform's (RAP) annual percent tree cover landsat satellite data (Robinson et al., 2019). Originally at 30-m spatial resolution but aggregated to 1 km. We used the cover value associated with the cell that a given camera station lies within. |
| Topographic Roughness | Continuous | Values represent the sum of the absolute value of the difference in elevation between each grid cell and its surrounding eight neighbors (3 x 3 window) (Kohl et al., 2019; Ruth et al., 2019). A value of 0 represents a flat surface while the maximum value represents the roughest terrain in the study area. Originally at 30-m spatial resolution but aggregated to 1 km. We used the roughness value associated with the cell that a given camera station lies within. |
| Snow Depth | Continuous | Daily snow depth data at 1-km grid cell accrued from SNODAS. We used the average value of all days within each session, so that snow depth values in a grid varied between but not within sessions. |
| Year | Categorical | Values represent the three winter sessions of our study (2021, 2022, 2023). |
| Detection Covariates |  |  |
| Snow Depth | Continuous | Daily snow depth data at 1-km grid cell accrued from SNODAS. We used the average value of all days within each weekly occasion, so that snow depth values at each station varied weekly. |
| Number of Cameras | Continuous, Range: 0-2 | Number of active/operational cameras at a given station. |
| Year | Categorical | Values represent the three winter sessions of our study (2021, 2022, 2023). |

**Table S2.** Detection and occupancy covariates in the best-supported first-order models for red fox, coyote, wolf, and cougar in each session.

| Session-Species | Detection model (p) | Top occupancy model ($\Psi$) |
| --- | --- | --- |
| Red Fox | *Year + Snow depth* | *Null (Year only)* |
| Coyote | *Null (Year only)* | *Null (Year only)* |
| Wolf | *Null (Year only)* | *Null (Year only)* |
| Cougar | *Year + Snow depth* | *Year + Snow depth + Roughness* |

**Table S3.** AIC model selection for all candidate first-order detection sub-models considered for each species. Bolded rows indicate selected models that were carried over into the occupancy model selection stage. K = number of model parameters, AIC = Akaike Information Criterion, ΔAIC = difference in AIC from the top model.

| First-order Detection Model | K | AIC | ΔAIC | AIC<sub>weight</sub> |
| --- | --- | --- | --- | --- |
| **Red Fox** |  |  |  |  |
| **p(Year + Snow depth) + $\Psi(.)$** | **11** | **7611** | **0** | **0.714** |
| p(Year + Active cameras + Snow depth) + $\Psi(.)$ | 12 | 7612.9 | 1.96 | 0.268 |
| p(Null (Year only)) + $\Psi(.)$ | 10 | 7619 | 8.01 | 0.013 |
| p(Year + Active cameras) + $\Psi(.)$ | 11 | 7620.8 | 9.86 | 0.005 |
| **Coyote** |  |  |  |  |
| **p(Null (Year only)) + $\Psi(.)$** | **10** | **7612.1** | **0** | **0.42** |
| p(Year + Active cameras) + $\Psi(.)$ | 11 | 7612.7 | 0.6 | 0.31 |
| p(Year + Snow depth) + $\Psi(.)$ | 11 | 7614.1 | 1.93 | 0.16 |
| p(Year + Active cameras + Snow depth) + $\Psi(.)$ | 12 | 7614.6 | 2.51 | 0.12 |
| **Wolf** |  |  |  |  |
| **p(Null (Year only)) + $\Psi(.)$** | **10** | **7616.6** | **0** | **0.39** |
| p(Year + Snow depth) + $\Psi(.)$ | 11 | 7617.5 | 0.91 | 0.25 |
| p(Year + Active cameras) + $\Psi(.)$ | 11 | 7617.8 | 1.17 | 0.22 |
| p(Year + Active cameras + Snow depth) + $\Psi(.)$ | 12 | 7618.7 | 2.08 | 0.14 |
| **Cougar** |  |  |  |  |
| p(Year + Active cameras + Snow depth) + $\Psi(.)$ | 12 | 7624.8 | 0 | 0.479 |
| **p(Year + Snow depth) + $\Psi(.)$** | **11** | **7625.1** | **0.23** | **0.427** |
| p(Null (Year only)) + $\Psi(.)$ | 10 | 7629.3 | 4.49 | 0.051 |
| p(Year + Active cameras) + $\Psi(.)$ | 11 | 7629.6 | 4.8 | 0.043 |

**Table S4.** AIC model selection for all candidate first-order occupancy models considered for each species. Each model includes the full detection sub-model. Bolded rows indicate selected models that were carried over into the occupancy model selection stage. K = number of model parameters, AIC = Akaike Information Criterion, ΔAIC = difference in AIC from the top model.

| First-order Detection + Occupancy Model | K | AIC | ΔAIC | AIC<sub>weight</sub> |
| --- | --- | --- | --- | --- |
| **Red Fox** |  |  |  |  |
| **p + $\Psi$(Null (Year only))** | **20** | **7588.1** | **0** | **0.222** |
| p + $\Psi$(Year + Roughness) | 21 | 7588.6 | 0.49 | 0.174 |
| p + $\Psi$(Year + Snow depth) | 21 | 7588.9 | 0.75 | 0.153 |
| p + $\Psi$(Year + Roughness + Snow depth) | 22 | 7589.2 | 1.01 | 0.134 |
| p + $\Psi$(Year + Cover) | 21 | 7589.6 | 1.48 | 0.106 |
| p + $\Psi$(Year + Roughness + Cover) | 22 | 7590 | 1.89 | 0.086 |
| p + $\Psi$(Year + Snow depth + Cover) | 22 | 7590.6 | 2.42 | 0.066 |
| p + $\Psi$(Year + Roughness + Snow depth + Cover) | 23 | 7590.8 | 2.66 | 0.059 |
| **Coyote** |  |  |  |  |
| p + $\Psi$(Year + Roughness + Cover) | 22 | 7587.3 | 0 | 0.215 |
| p + $\Psi$(Year + Cover) | 21 | 7587.6 | 0.24 | 0.190 |
| p + $\Psi$(Year + Roughness) | 21 | 7588.2 | 0.84 | 0.141 |
| **p + $\Psi$(Null (Year only))** | **20** | **7588.7** | **1.38** | **0.108** |
| p + $\Psi$(Year + Snow depth + Cover) | 22 | 7588.8 | 1.41 | 0.106 |
| p + $\Psi$(Year + Roughness + Snow depth + Cover) | 23 | 7588.9 | 1.53 | 0.100 |
| p + $\Psi$(Year + Roughness + Snow depth) | 22 | 7589.5 | 2.19 | 0.072 |
| p + $\Psi$(Year + Snow depth) | 21 | 7589.6 | 2.27 | 0.069 |
| **Wolf** |  |  |  |  |
| **p + $\Psi$(Null (Year only))** | **20** | **7588.9** | **0** | **0.221** |
| p + $\Psi$(Year + Snow depth) | 21 | 7589.4 | 0.59 | 0.165 |
| p + $\Psi$(Year + Roughness + Snow depth) | 22 | 7589.6 | 0.71 | 0.155 |
| p + $\Psi$(Year + Roughness) | 21 | 7589.6 | 0.74 | 0.153 |
| p + $\Psi$(Year + Cover) | 21 | 7590.6 | 1.71 | 0.094 |
| p + $\Psi$(Year + Snow depth + Cover) | 22 | 7590.9 | 2.08 | 0.078 |
| p + $\Psi$(Year + Roughness + Snow depth + Cover) | 23 | 7591.1 | 2.27 | 0.071 |
| p + $\Psi$(Year + Roughness + Cover) | 22 | 7591.4 | 2.52 | 0.063 |
| **Cougar** |  |  |  |  |
| p + $\Psi$(Year + Roughness + Snow depth + Cover) | 23 | 7542.4 | 0 | 0.54 |
| **p + $\Psi$(Year + Roughness + Snow depth)** | **22** | **7543.8** | **1.35** | **0.28** |
| p + $\Psi$(Year + Roughness + Cover) | 22 | 7545.6 | 3.15 | 0.11 |
| p + $\Psi$(Year + Roughness) | 21 | 7546.6 | 4.22 | 0.066 |
| p + $\Psi$(Year + Snow depth) | 21 | 7578.8 | 36.39 | 6.8e-9 |
| p + $\Psi$(Year + Snow depth + Cover) | 22 | 7579.4 | 36.97 | 5.1e-9 |
| p + $\Psi$(Null (Year only)) | 20 | 7586.2 | 43.79 | 1.7e-10 |
| p + $\Psi$(Year + Cover) | 21 | 7587.3 | 44.86 | 9.9e-11 |

**Table S5.**
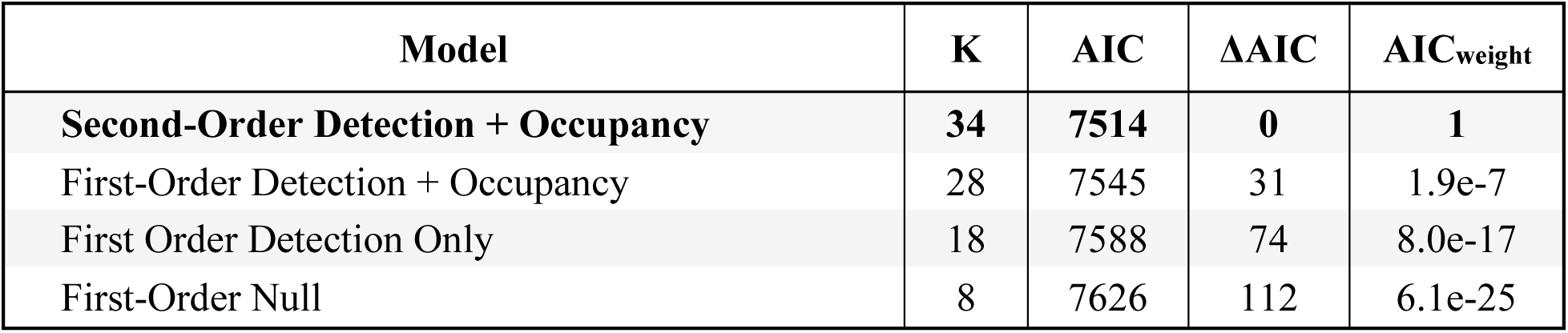
AIC model comparison for all candidate first- and second-order multi-species occupancy models considered. Bolded rows indicate selected models that we reported results for. K = number of model parameters, AIC = Akaike Information Criterion, ΔAIC = difference in AIC from the top model.

| Model | K | AIC | ΔAIC | AIC<sub>weight</sub> |
| --- | --- | --- | --- | --- |
| **Second-Order Detection + Occupancy** | **34** | **7514** | **0** | **1** |
| First-Order Detection + Occupancy | 28 | 7545 | 31 | 1.9e-7 |
| First Order Detection Only | 18 | 7588 | 74 | 8.0e-17 |
| First-Order Null | 8 | 7626 | 112 | 6.1e-25 |

**Table S6.**
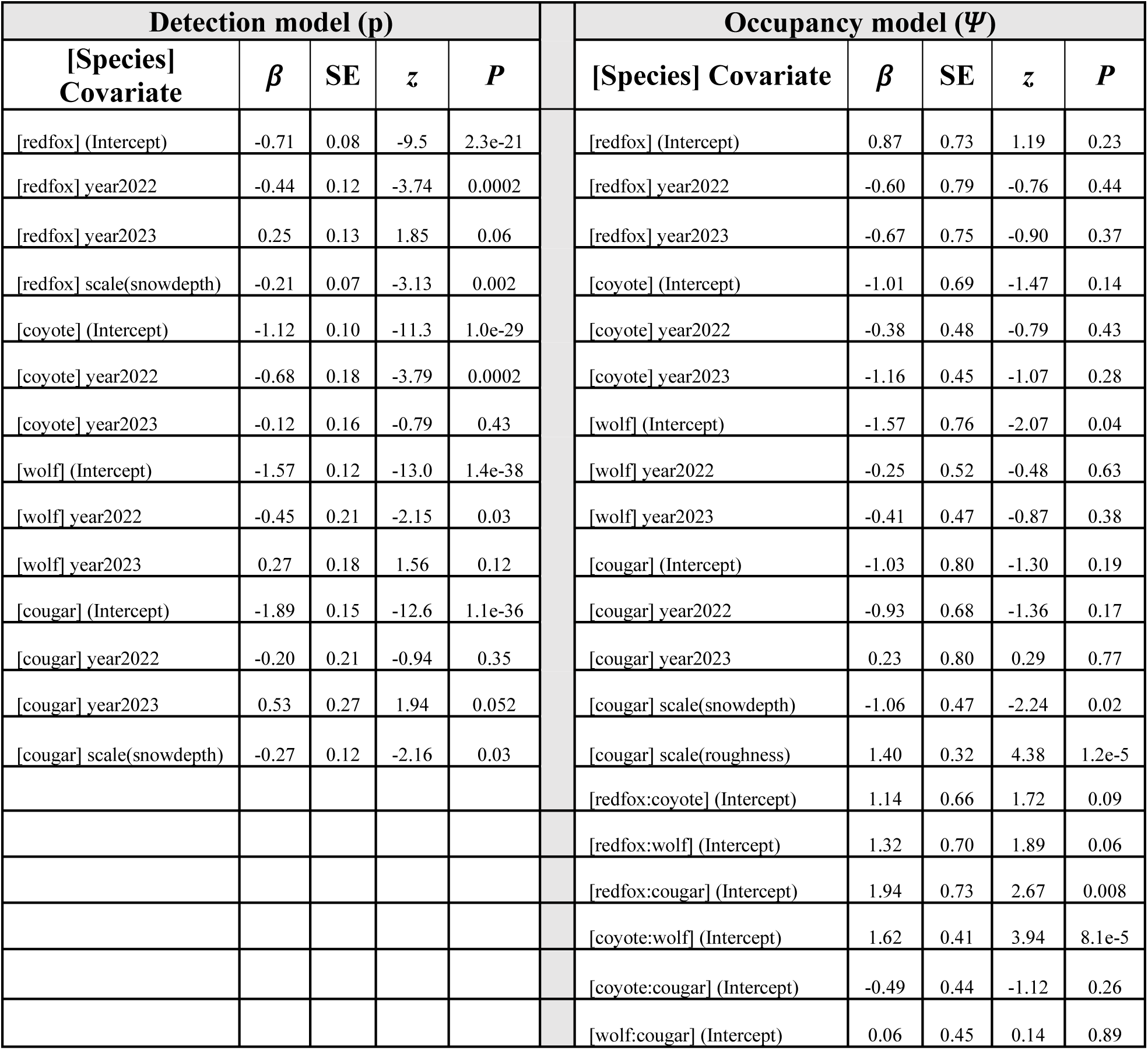
Beta coefficient estimates for detection and occupancy covariates retained in the top second-order multi-species occupancy model for each session.

**Figure S1.**
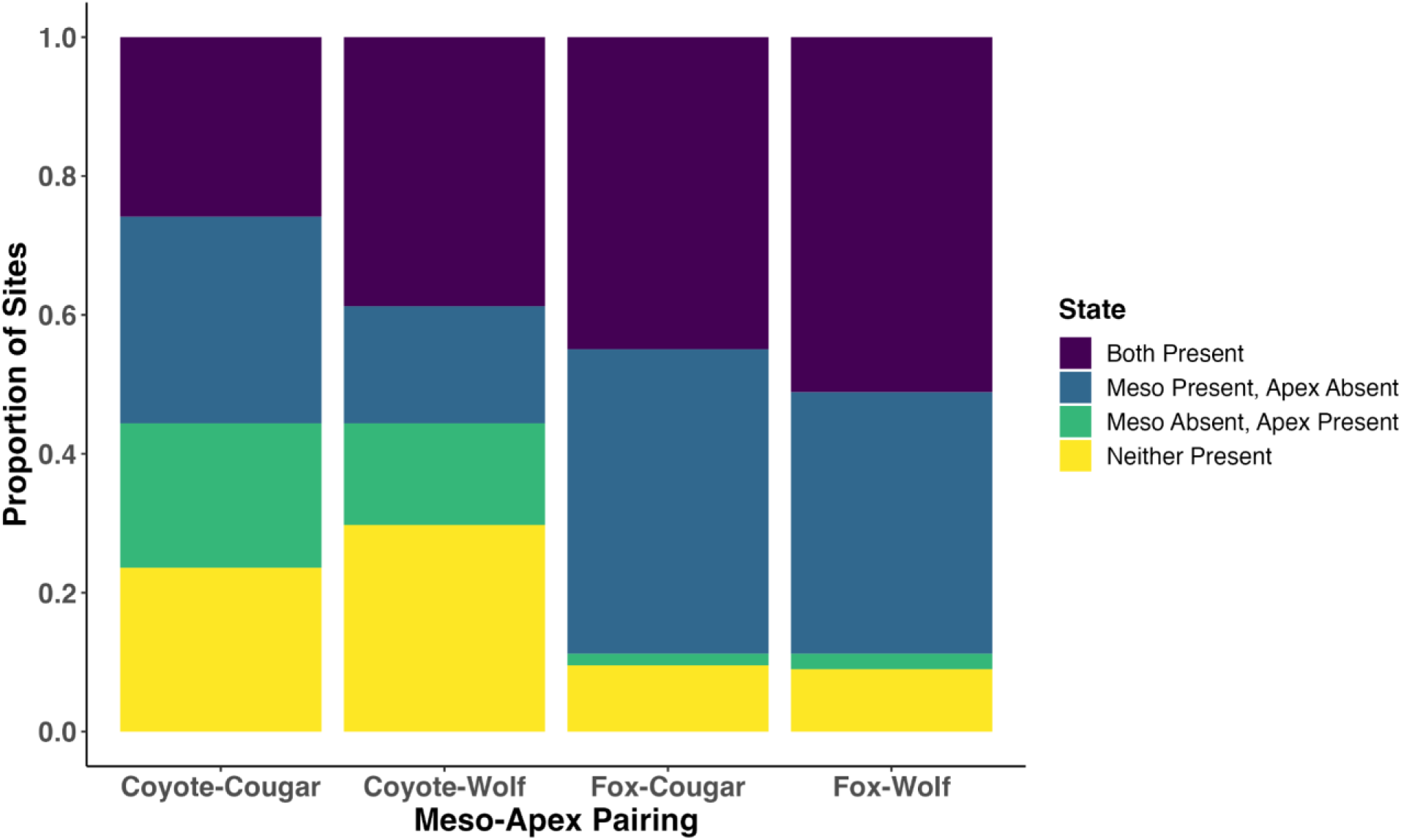
Proportion of camera sites exhibiting each co-occurrence state (*z*) for each meso-apex species pair.

**Figure S2.**
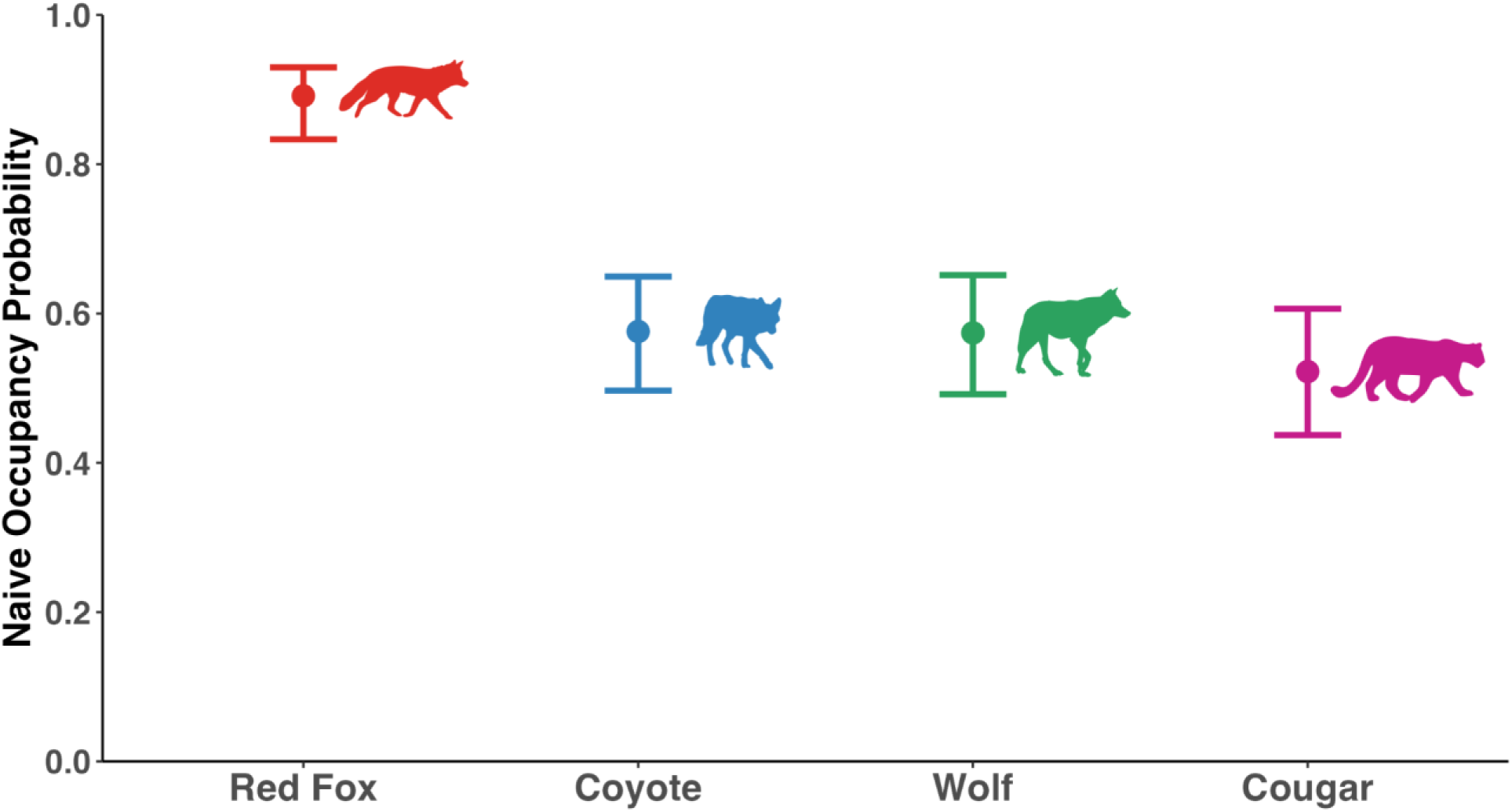
Naive occupancy probability (proportion of sites occupied) across all sites for each species.

**Figure S3.**
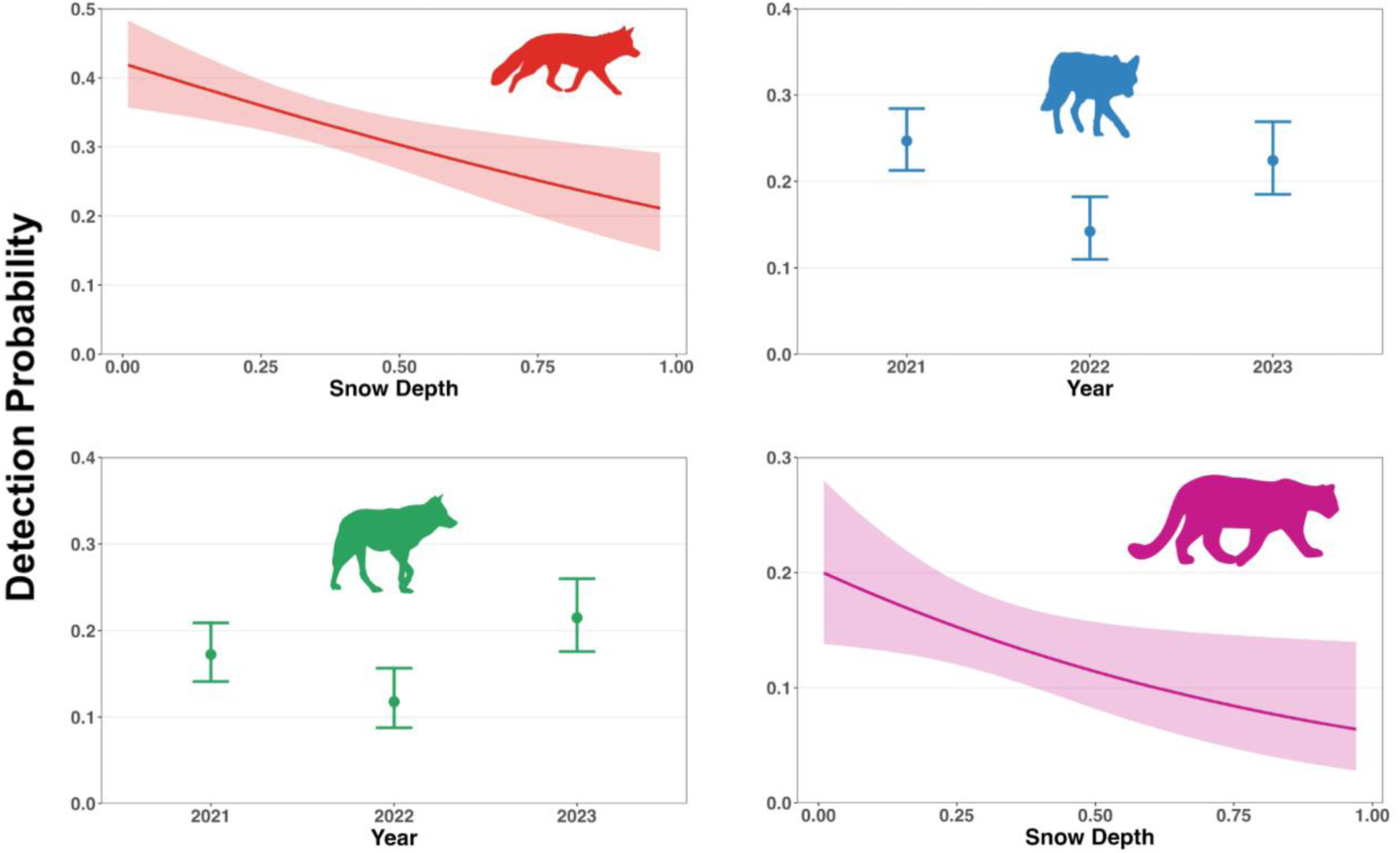
Detection probabilities for each species as a function of significant covariates retained in the top second-order multi-species occupancy model. Shaded regions are 95% confidence intervals.

**Figure S4.**
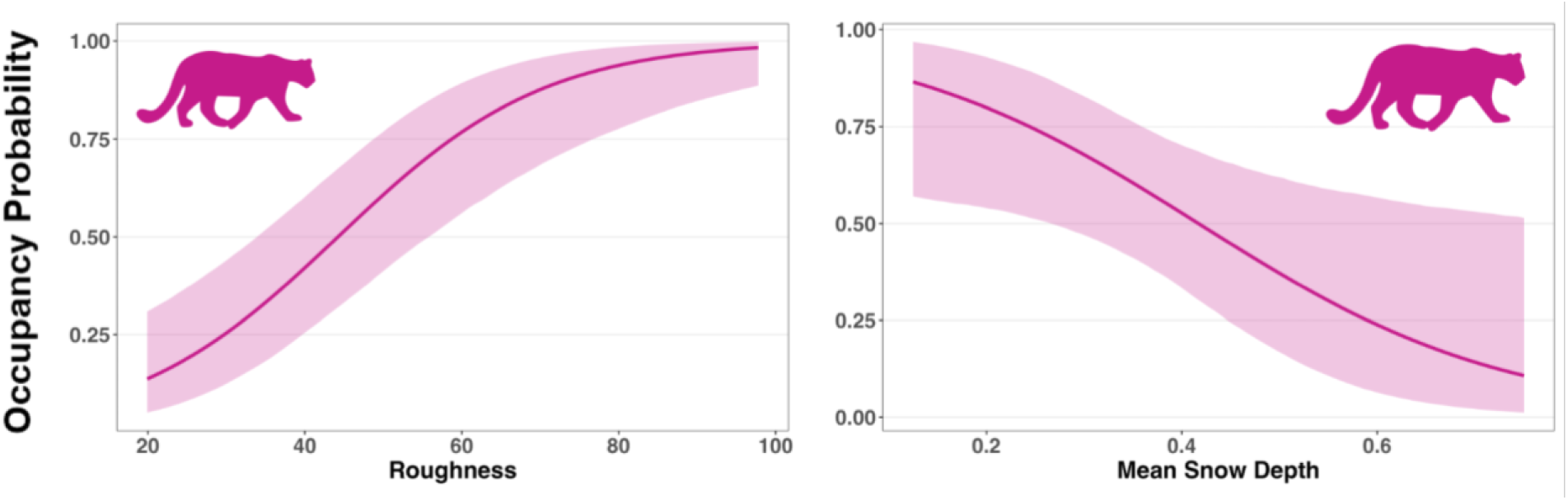
Marginal occupancy probability for cougar as a function of habitat covariates retained in the top second-order multi-species occupancy model. Shaded regions are 95% confidence intervals.

**Figure S5.**
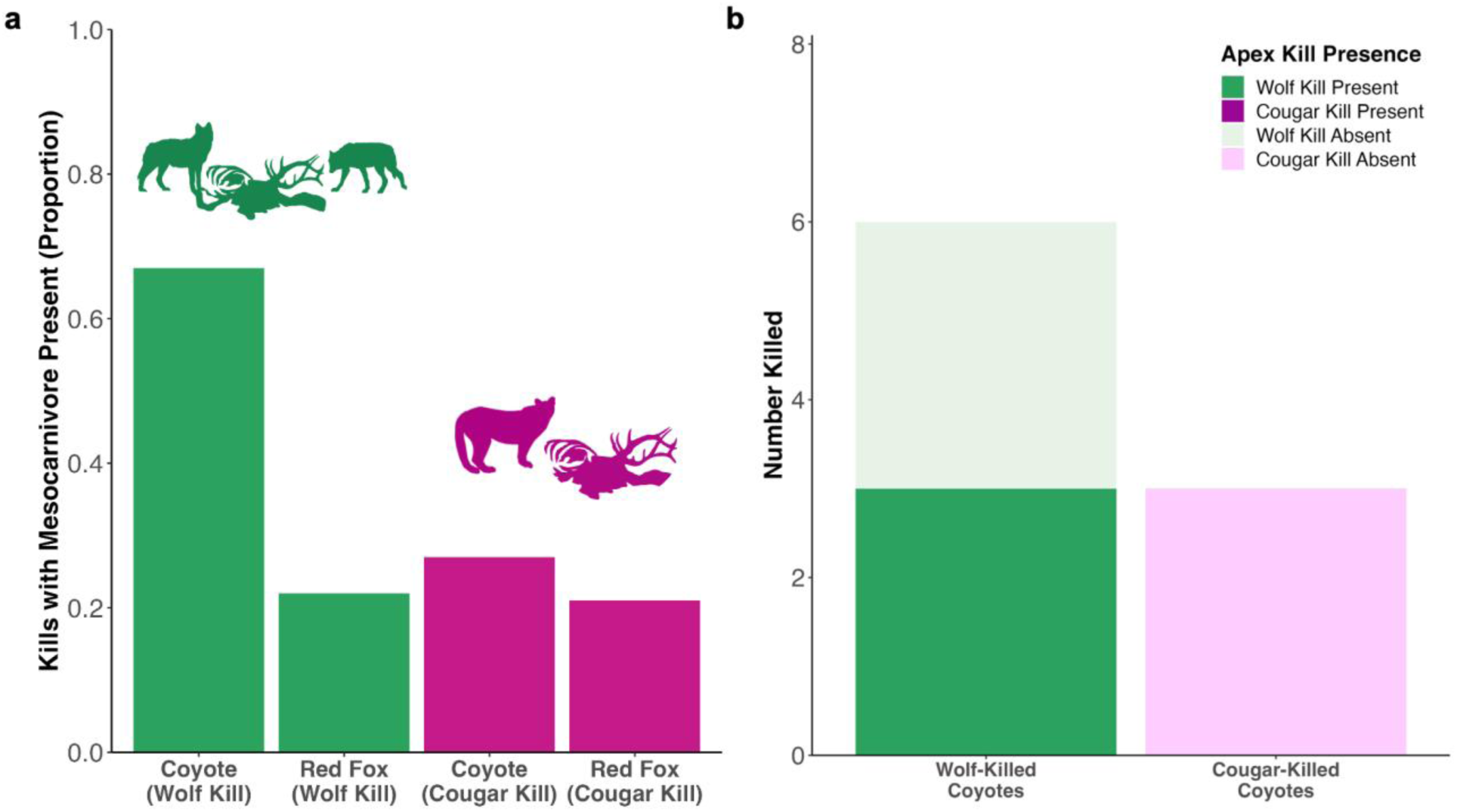
Wolf (n = 113) and cougar (n = 67) kills, surveyed through GPS-cluster searches of collared wolves and cougars during our three camera-trap sessions (2021-2023), bearing **a)** presence of coyote and red fox scavenging expressed as a proportion, and **b**) counts of coyote and red fox killed. For methods see main text. Wolf and elk carcass graphics by Kira Cassidy.

## REFERENCES

1. Ripple WJ et al. 2014 Status and Ecological Effects of the World’s Largest Carnivores. Science 343, 1241484. (doi:10.1126/science.1241484)

2. Bangs EE, Fritts SH. 1996 The Reintroduction of Gray Wolves to Yellowstone National Park and Central Idaho - Final Environmental Impact Statement 1994. Wildl. Soc. Bull. , 414.

3. Chapron G et al. 2014 Recovery of large carnivores in Europe’s modern human-dominated landscapes. Science 346, 1517–1519. (doi:10.1126/science.1257553)

4. Lotze HK, Coll M, Magera AM, Ward-Paige C, Airoldi L. 2011 Recovery of marine animal populations and ecosystems. Trends Ecol. Evol. 26, 595–605. (doi:10.1016/j.tree.2011.07.008)

5. Levi T, Wilmers CC. 2012 Wolves–coyotes–foxes: a cascade among carnivores. Ecology 93, 921–929. (doi:10.1890/11-0165.1)

6. Brunet MJ et al. 2022 Cats and dogs: A mesopredator navigating risk and reward provisioned by an apex predator. Ecol. Evol. 12, e8641. (doi:10.1002/ece3.8641)

7. Sivy KJ, Pozzanghera CB, Grace JB, Prugh LR. 2017 Fatal Attraction? Intraguild Facilitation and Suppression among Predators. Am. Nat. 190, 663–679. (doi:10.1086/693996)

8. Wilmers CC, Crabtree RL, Smith DW, Murphy KM, Getz WM. 2003 Trophic facilitation by introduced top predators: grey wolf subsidies to scavengers in Yellowstone National Park. J. Anim. Ecol. 72, 909–916. (doi:10.1046/j.1365-2656.2003.00766.x)

9. Ruprecht J et al. 2021 Variable strategies to solve risk–reward tradeoffs in carnivore communities. Proc. Natl. Acad. Sci. 118, e2101614118. (doi:10.1073/pnas.2101614118)

10. Prugh LR, Sivy KJ. 2020 Enemies with benefits: integrating positive and negative interactions among terrestrial carnivores. Ecol. Lett. 23, 902–918. (doi:10.1111/ele.13489)

11. Broekhuis F. 2015 Cat eats cat: leopard consumes adult cheetah, Maasai Mara Game Reserve, Kenya. CAT News 65, 33–34.

12. Houston DB. 1982 The Northern Yellowstone Elk: Ecology and Management. Macmillan.

13. White PJ, Wallen RL, Hallac D, Auttelet K, Jerret J. 2015 Yellowstone Bison: Conserving an American Icon in Modern Society. Yellowstone Association.

14. Smith DW, Stahler DR, MacNulty DR. 2020 Yellowstone Wolves: Science and Discovery in the World’s First National Park. University of Chicago Press.

15. Metz MC, Smith DW, Vucetich JA, Stahler DR, Peterson RO. 2012 Seasonal patterns of predation for gray wolves in the multi-prey system of Yellowstone National Park. J. Anim. Ecol. 81, 553–563. (doi:10.1111/j.1365-2656.2011.01945.x)

16. Metz MC. 2021 Estimating wolf predation metrics, patterns, and dynamics across time and space in the multi-prey system of Yellowstone National Park. University of Montana. See https://scholarworks.umt.edu/etd/11842/.

17. Ruth T, Buotte P, Hornocker M. 2019 Yellowstone Cougars: Ecology Before And During Wolf Restoration. University Press of Colorado. (doi:10.5876/9781607328292)

18. Plumb GE, White PJ, Coughenour MB, Wallen RL. 2009 Carrying capacity, migration, and dispersal in Yellowstone bison. Biol. Conserv. 142, 2377–2387. (doi:10.1016/j.biocon.2009.05.019)

19. Sun CC, Fuller AK, Royle JA. 2014 Trap Configuration and Spacing Influences Parameter Estimates in Spatial Capture-Recapture Models. PLOS ONE 9, e88025. (doi:10.1371/journal.pone.0088025)

20. Anton C. 2020 The Demography and Comparative Ethology of Top Predators in a Multi-Carnivore System. UC Santa Cruz. See https://escholarship.org/uc/item/3xd9379b.

21. Greenberg S, Godin T, Whittington J. 2019 Design patterns for wildlife-related camera trap image analysis. Ecol. Evol. 9, 13706–13730. (doi:10.1002/ece3.5767)

22. Rota CT, Ferreira MAR, Kays RW, Forrester TD, Kalies EL, McShea WJ, Parsons AW, Millspaugh JJ. 2016 A multispecies occupancy model for two or more interacting species. Methods Ecol. Evol. 7, 1164–1173. (doi:10.1111/2041-210X.12587)

23. Fiske I, Chandler R. 2011 **unmarked** : An *R* Package for Fitting Hierarchical Models of Wildlife Occurrence and Abundance. J. Stat. Softw. 43. (doi:10.18637/jss.v043.i10)

24. R Core Team. 2023 R: A Language and Environment for Statistical Computing.

25. MacKenzie DI, editor. 2006 Occupancy estimation and modeling: inferring patterns and dynamics of species. Amsterdam ; Boston: Elsevier.

26. Steenweg R, Hebblewhite M, Whittington J, Lukacs P, McKelvey K. 2018 Sampling scales define occupancy and underlying occupancy–abundance relationships in animals. Ecology 99, 172–183. (doi:10.1002/ecy.2054)

27. National Operational Hydrologic Remote Sensing Center. 2004 Snow Data Assimulation System (SNODAS) Data Products at NSIDC. (10.7265/N5TB14TC)

28. Cano-Martínez R, Thorsen NH, Hofmeester TR, Odden J, Linnell J, Devineau O, Angoh SYJ, Odden M. 2024 Bottom-up rather than top-down mechanisms determine mesocarnivore interactions in Norway. Ecol. Evol. 14, e11064. (doi:10.1002/ece3.11064)

29. Kéry M, Royle JA. 2016 Distribution, Abundance, and Species Richness in Ecology. Elsevier. (doi:10.1016/B978-0-12-801378-6.00001-1)

30. Kéry M, Royle JA. 2021 Applied Hierarchical Modeling in Ecology: Analysis of Distribution, Abundance and Species Richness in R and BUGS. Elsevier. (doi:10.1016/B978-0-12-809585-0.00001-6)

31. Twining JP, Brazeal JL, Jensen PG, Fuller AK. 2024 Intraguild interactions and abiotic conditions mediate occupancy of mammalian carnivores: co-occurrence of coyotes–fishers– martens. Oikos 2024, e10577. (doi:10.1111/oik.10577)

32. Arnold TW. 2010 Uninformative Parameters and Model Selection Using Akaike’s Information Criterion. J. Wildl. Manag. 74, 1175–1178. (doi:10.1111/j.1937-2817.2010.tb01236.x)

33. Rowcliffe M. 2023 **activity**: Animal Activity Statistics. 1.3.4 (url:10.32614/CRAN.package.activity)

34. Tanwar KS, Sadhu A, Jhala YV. 2021 Camera trap placement for evaluating species richness, abundance, and activity. Sci. Rep. 11, 23050. (doi:10.1038/s41598-021-02459-w)

35. Ridout MS, Linkie M. 2009 Estimating overlap of daily activity patterns from camera trap data. J. Agric. Biol. Environ. Stat. 14, 322–337. (doi:10.1198/jabes.2009.08038)

36. Cusack JJ, Dickman AJ, Kalyahe M, Rowcliffe JM, Carbone C, MacDonald DW, Coulson T. 2017 Revealing kleptoparasitic and predatory tendencies in an African mammal community using camera traps: a comparison of spatiotemporal approaches. Oikos 126, 812–822. (doi:10.1111/oik.03403)

37. Davis RS, Yarnell RW, Gentle LK, Uzal A, Mgoola WO, Stone EL. 2021 Prey availability and intraguild competition regulate the spatiotemporal dynamics of a modified large carnivore guild. Ecol. Evol. 11, 7890–7904. (doi:10.1002/ece3.7620)

38. Anderson CR, Lindzey FG. 2003 Estimating Cougar Predation Rates from GPS Location Clusters. J. Wildl. Manag. 67, 307. (doi:10.2307/3802772)

39. Perrig PL, Lambertucci SA, Donadio E, Smith JA, Middleton AD, Pauli JN. 2023 Risk effects cascade up to an obligate scavenger. Ecology 104, e3871. (doi:10.1002/ecy.3871)

40. Smith AB, Squires JR, Bjornlie NL, Holbrook JD. 2023 Divergent or convergent: how do forest carnivores use time in the Greater Yellowstone Ecosystem? J. Mammal., gyad070. (doi:10.1093/jmammal/gyad070)

41. Fedriani JM, Fuller TK, Sauvajot RM, York EC. 2000 Competition and intraguild predation among three sympatric carnivores. Oecologia 125, 258–270. (doi:10.1007/s004420000448)

42. Van Etten KW, Wilson KR, Crabtree RL. 2007 Habitat Use of Red Foxes in Yellowstone National Park Based on Snow Tracking and Telemetry. J. Mammal. 88, 1498–1507. (doi:10.1644/07-MAMM-A-076.1)

43. Jensen AJ, Marneweck CJ, Kilgo JC, Jachowski DS. 2022 Coyote diet in North America: geographic and ecological patterns during range expansion. Mammal Rev. 52, 480–496. (doi:10.1111/mam.12299)

44. Allen ML, Elbroch LM, Wilmers CC, Wittmer HU. 2014 Trophic Facilitation or Limitation? Comparative Effects of Pumas and Black Bears on the Scavenger Community. PLoS ONE 9, e102257. (doi:10.1371/journal.pone.0102257)

45. Berger KM, Gese EM. 2007 Does interference competition with wolves limit the distribution and abundance of coyotes? J. Anim. Ecol. 76, 1075–1085. (doi:10.1111/j.1365-2656.2007.01287.x)

46. Merkle JA, Stahler DR, Smith DW. 2009 Interference competition between gray wolves and coyotes in Yellowstone National Park. Can. J. Zool. 87, 56–63. (doi:10.1139/Z08-136)

47. Sinclair ARE. 2003 Mammal population regulation, keystone processes and ecosystem dynamics. Philos. Trans. R. Soc. Lond. B. Biol. Sci. 358, 1729–1740. (doi:10.1098/rstb.2003.1359)

48. Brice EM, Larsen EJ, MacNulty DR. 2022 Sampling bias exaggerates a textbook example of a trophic cascade. Ecol. Lett. 25, 177–188. (doi:10.1111/ele.13915)

49. Hobbs NT, Johnston DB, Marshall KN, Wolf EC, Cooper DJ. 2024 Does restoring apex predators to food webs restore ecosystems? Large carnivores in Y ellowstone as a model system. Ecol. Monogr. 94, e1598. (doi:10.1002/ecm.1598)

50. Ripple WJ, Beschta RL. 2012 Trophic cascades in Yellowstone: The first 15years after wolf reintroduction. Biol. Conserv. 145, 205–213. (doi:10.1016/j.biocon.2011.11.005)

## REFERENCES

Kohl, M. T., Ruth, T. K., Metz, M. C., Stahler, D. R., Smith, D. W., White, P. J., & MacNulty, D. R. (2019). Do prey select for vacant hunting domains to minimize a multi-predator threat? Ecology Letters, 22(11), 1724–1733. 10.1111/ele.13319

Robinson, N. P., Jones, M. O., Moreno, A., Erickson, T. A., Naugle, D. E., & Allred, B. W. (2019). Rangeland Productivity Partitioned to Sub-Pixel Plant Functional Types. Remote Sensing, 11(12), 1427. 10.3390/rs11121427

Ruth, T., Buotte, P., & Hornocker, M. (2019). Yellowstone Cougars: Ecology Before And During Wolf Restoration. University Press of Colorado. 10.5876/9781607328292

